# Using gene genealogies to localize rare variants associated with complex traits in diploid populations

**DOI:** 10.1101/182345

**Authors:** Charith B. Karunarathna, Jinko Graham

**Affiliations:** Department of Statistics and Actuarial Science, Simon Fraser University, Burnaby, BC, Canada.

## Abstract

**Background and Aims:** Many methods can detect trait association with causal variants in candidate genomic regions; however, a comparison of their ability to localize causal variants is lacking. We extend a previous study of the detection abilities of these methods to a comparison of their localization abilities.

**Methods:** Through coalescent simulation, we compare several popular association methods. Cases and controls are sampled from a diploid population to mimic human studies. As benchmarks for comparison, we include two methods that cluster phenotypes on the true genealogical trees, a naive Mantel test considered previously in haploid populations and an extension that takes into account whether case haplotypes carry a causal variant. We first work through a simulated dataset to illustrate the methods. We then perform a simulation study to score the localization and detection properties.

**Results:** In our simulations, the association signal was localized least precisely by the naive Mantel test and most precisely by its extension. Most other approaches had intermediate performance similar to the single-variant Fisher’s-exact test.

**Conclusions:** Our results confirm earlier findings in haploid populations about potential gains in performance from genealogy-based approaches. They also highlight differences between haploid and diploid populations when localizing and detecting causal variants.

## 1 Introduction

Most genetic association studies focus on common variants, but rare variants can play major roles in influencing complex traits [1, 2]. Rare causal variants identified through sequencing could thus explain some of the missing heritability of complex traits [3]. However, for rare variants, standard methods to test for association with single genetic variants are underpowered unless sample sizes are very large [4]. The lack of power of single-variant approaches holds in fine-mapping as well as genome-wide association studies.

In this report, we are concerned with fine-mapping a candidate genomic region that has been sequenced in cases and controls to identify a disease-risk locus. Our work extends an earlier comparison of methods for *detecting* disease association in a candidate genomic region [5] to a comparison of methods for *localizing* the association signal. Additionally, in the current investigation, we sample cases and controls from a diploid or two-parent population to mimic studies in humans. In the previous investigation, cases and controls were sampled from a haploid or one-parent population.

A number of methods have been developed to evaluate the disease association for both a single variant and multiple variants in a genomic region. Besides single-variant methods, we consider three broad classes of methods for analysing association: pooled-variant, joint-modelling and tree-based methods. Pooled-variant methods evaluate the cumulative effects of multiple genetic variants in a genomic region. The score statistics from marginal models of the trait association with individual variants are collapsed into a single test statistic by combining the information for multiple variants into a single genetic score [4]. Joint-modeling methods model the joint effect of multiple genetic variants on the trait simultaneously. These methods can assess whether a variant carries any further information about the trait beyond what is explained by the other variants. When trait-influencing variants are in low linkage disequilibrium, this approach may be more powerful than pooling test statistics for marginal associations across variants [6]. Tree-based methods assess whether trait values co-cluster with the local genealogical tree for the haplotypes (e.g., [7, 8]). A local genealogical tree represents the ancestry of the sample of haplotypes at each locus. Haplotypes carrying the same causal alleles are expected to be related and cluster on the genealogical tree at a disease-risk locus.

In practice, true trees are unknown. However, clustering statistics based on true trees represent a best case for detecting or localizing association because tree uncertainty is eliminated. Burkett et al. [5] used known trees to assess the effectiveness of tree-based approaches for detection of disease-risk variants in a haploid population. They found that clustering statistics computed on the known trees outperform popular methods for detecting causal variants in a candidate genomic region. Following Burkett et al. [5], we use Mantel tests that associate phenotypic and genealogical distances as the clustering statistics. These statistics, which rely on known trees, serve as benchmarks against which to compare the popular association methods. However, unlike Burkett et al. [5], who focus on detection of disease-risk variants, we focus on localization of association signal in the candidate genomic region. Additionally, we use a diploid rather than a haploid disease model to mimic human populations.

In this report, we compare the ability of several popular methods of association mapping to localize causal variants in a subregion of a larger, candidate, genomic region. In our simulation study, we use sequence data generated under an approximation to the coalescent with recombination [9]. To illustrate ideas, we start by working through a particular example dataset as a case study for insight into the association methods. We next perform a simulation study involving 200 sequencing datasets and score which association method localizes the risk subregion most precisely. We conclude with a summary and discussion of our results. Our results confirm the earlier findings [5] indicating potential gains in performance from ancestral tree-based approaches. They also highlight some important differences between haploid and diploid populations when localizing causal variants.

## 2 Methods

In this section, we describe our data simulation, the association methods we considered and the way we assessed localization and detection of the association signal. We next describe the popular association methods we evaluated for fine mapping. We finally explain the simulation study involving 200 sequencing datasets to address the signal localization and detection.

### 2.1 Data simulation

First, we report how we simulated haplotype data from ancestral trees. Second, we describe how we assigned the disease status to individuals and sampled data for our case-control study. We used fastsimcoal2 [10] to simulate ancestral trees and 3000 haplotypes of 4000 equispaced single-nucleotide variants (SNVs) in a 2 million base-pair (Mbp) genomic region. We used a recombination rate of 1 × 10^−8^ per base-pair per generation [11], in a diploid population of constant effective size, *N_e_* = 6200 [12]. To mimic random mating in a diploid population, we then randomly paired the 3000 haplotypes into 1500 diploid individuals. Next, disease status was assigned to the 1500 individuals based on randomly-sampled risk SNVs from the middle genomic region of 950 — 1050 kbp. For risk SNVs, the number of copies of the derived (i.e. mutant) allele increased the risk of disease according to a logistic regression model,

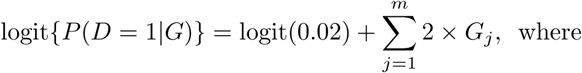

- logit(*p*) = log[*p*/(1 − *p*)] for 0 < *p* < 1,
- *D* is disease status (*D* = 1, case; *D* = 0, control),
- *G* = (*G*_1_, *G*_2_,…, *G_m_*) is an individual’s multi-locus genotype at *m* risk SNVs, with *G_j_* being the number of copies of the derived allele at the *j^th^* risk SNV, and
- the value of the intercept term is chosen to ensure that the probability of sporadic disease (i.e. 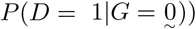 is approximately 2%.

To select risk SNVs in the model, we randomly sampled SNVs from the middle subregion one-at-a-time, until the disease prevalence was between 9.5 — 10.5% in the 1500 individuals. Our selection of risk SNVs is not restricted by the minor allele frequency and therefore differs slightly from Burkett et al. [5], which allowed only SNVs with MAF < 1% to be risk SNVs. After assigning disease status to the 1500 individuals, we randomly sampled 50 cases (i.e. diseased) from the affected individuals and 50 controls (i.e. non-diseased) from the unaffected individuals. We then extracted the data for the variable SNVs in the resulting case-control sample to examine the patterns of disease association in subsequent analyses.

### 2.2 Association analysis

In this section, we review the methods for association mapping that we considered. These methods fall under four categories: single-variant method, pooled-variant methods, joint-modeling methods and tree-based methods.

#### 2.2.1 Single-variant method

For the single-variant method, we used the standard Fisher’s exact test of disease association with each of the SNVs in the case-control sample. In Fisher’s exact test, each of the variant sites in the case-control sample was tested for an association with the disease outcome using a 2 × 3 table to compare genotype frequencies. This single-variant association was assessed with the p-value of Fisher’s exact test on the contingency table. Each row in a table represents disease status of individuals, and a column represents the three possible genotypes. The *−log_10_* p-value from the test was recorded as the association signal for each variant. However, single-variant tests are less powerful for rare variants than for common variants [13]. We therefore considered three other ways to assess the association signal based on pooled-variant, joint-modelling and tree-based methods.

#### 2.2.2 Pooled-variant methods

For the pooled-variant methods, we evaluated the Variable Threshold (VT) and the C-alpha test. The variable threshold approach of Price et al. [14], use a generalized linear model to relate the phenotypes to the counts of variants in the genomic region of interest which have minor allele frequencies (MAFs) below some user-defined threshold (e.g. 1% or 5%). The idea is that variants with MAF below the threshold have a higher prior probability of being functional than the variants with higher MAF, based on population-genetic arguments. For each possible MAF threshold, VT computes a score measuring the strength of association between the pheonotype and the genomic region, and uses the maximum of the score over all allele frequency thresholds. The statistical significance of the maximum score is then assessed by a permutation test. Price et al. [14] found that the VT approach had high power to detect the association between rare variants and disease traits when effects are in one direction. Unlike the VT test, the C-alpha test of Neale et al. [15] is a variance-components approach that assumes the effects of variants are random with mean zero. The C-alpha procedure tests the variance of genetic effects under the assumption that variants observed in cases and controls are a mixture of risk, protective or neutral variants. Neale et al. [15] found that the C-alpha test showed greater power than burden tests such as VT when the effects are bi-directional. We applied the VTWOD function in the R package RVtests [16] for the VT-test and the SKAT function in the R package SKAT [17] for the C-alpha test. We used sliding windows of 20 SNVs overlapping by 5 SNVs across the simulated region.

#### 2.2.3 Joint-modeling methods

For the joint-modeling methods, we evaluated the CAVIARBF [18] and elastic-net [19] methods. CAVIARBF is a fine-mapping method that uses marginal test statistics for the SNVs and their pairwise association to approximate the Bayesian multiple regression of phenotypes onto variants that is implemented in BIMBAM [20]. However, CAVIARBF is much faster than BIMBAM because it computes Bayes factors using only the SNVs in each causal model rather than all SNVs. These Bayes factors can be used to calculate the posterior probability of SNVs in the region being causal; i.e. the posterior inclusion probability (PIP). To compute PIPs for SNVs, a set of models and their Bayes factors have to be considered. Let *p* be the total number of SNVs in a candidate region; then the number of possible causal models is 2^*p*^. To reduce the number of causal models to evaluate and thus save computational time and effort, CAVIARBF imposes a limit, *L*, on the number of causal variants. This limitation reduces the number of models in the model space from 2^*p*^ to 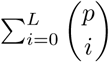. Since there were 2747 SNVs in our example dataset, to keep the computational load down, we considered *L* = 2 throughout this investigation.

The elastic net [19] is a hybrid regularization and variable selection method that linearly combines the L1 and L2 regularization penalties of the lasso [21] and ridge (e.g., [22]) regression methods in multiple regression. This combination of lasso and ridge penalties provides a more precise prediction than using multiple regression, when SNVs are in high linkage disequilibrium [23]. In addition, the elastic net can accommodate situations in which the number of predictors exceeds the number of observations. We used the elastic net to select risk SNVs by considering only the main effects. The variable inclusion probability (VIP), a frequentist analog of the Bayesian posterior inclusion probability was used as a measure of the importance of a SNV for predicting disease risk [6]. To obtain the VIP for a SNV, we re-fitted the elastic-net model using 100 bootstrap samples and calculated the proportion of samples in which the SNV was included in the fitted model. In our analysis, we applied the elastic net using the R package glmnet [24].

#### 2.2.4 Tree-based methods

We considered two tree-based methods to assess clustering of disease status on the gene genealogy connecting haplotypes at a putative risk variant: Blossoc (BLOck aSSOCiation; [8]), which uses reconstructed trees, and a Mantel test which uses the true trees. Blossoc aims to localize the risk variants by reconstructing genealogical trees at each SNV. The reconstructed trees approximate perfect phylogenies [25] for each SNV, assuming an infinite-sites model of mutation. These trees are scored according to the non-random clustering of affected individuals. The underlying idea is that genomic regions containing SNVs with high clustering scores are likely to harbour risk variants. Blossoc can be used for both phased and unphased genotype data. However, the method is impractical to apply to unphased data with more than a few SNVs due to the computational burden associated with phasing. We therefore assumed the SNV data are phased, as might be done in advance with a fast-phasing algorithm such as fastPHASE [26], BEAGLE [27], IMPUTE2 [28] or MACH [29, 30]. We evaluated Blossoc with the phased haplotypes, using the probability-score criterion which is the recommended scoring scheme for small datasets [8].

In practice, the true trees are unknown but as the data were simulated we had access to this information. Also, the cluster statistics based on true trees represent a best case insofar as tree uncertainty is eliminated [5]. We therefore included two versions of the Mantel test as a benchmark for comparison. In the first version, the phenotype corresponding to a haplotype is scored according to whether or not the haplotype comes from a case. We refer to the first version as the *naive* Mantel test because all case haplotypes are treated the same, even those not carrying any risk variants. In the second version, the phenotype is scored according to whether or not the haplotype comes from a case and carries a risk variant. We refer to the second version as the *informed* Mantel test because it takes into account whether or not a case haplotype carries a risk variant. The informed Mantel test is a best-case scenario insofar as the uncertainty about the risk-variant-carrying status of the case haplotypes is eliminated. Both Mantel tests correlate the pairwise distance in the known ancestry with those in the phenotypes. Following Burkett et al. [5], we used pairwise distances calculated from the rank of the coalescent event rather than the actual times on the tree. To focus on the rare variants, the test statistic upweights the short branches close to present at the tip of the tree, by assigning a branch-length of one to all branches, even the relatively longer branches that are expected to occur close to the time to the most recent common ancestor. Pairwise distances between haplotypes on this re-scaled tree are then correlated to pairwise phenotypic distances. We determined the distance measures, *d_ij_* = 1 − *s_ij_*, where *s_ij_* = (*y_i_* − *μ*)(*y_j_* − *μ*) is the similarity score between haplotype *i* and *j*, *y_i_* is the binary phenotype (coded as 0 or 1) and *μ* is the disease prevalence in the 1500 simulated individuals. We then used the Mantel statistic to compare the phenotype-based distance matrix, *d*, with the re-scaled tree-distance matrix. Note that we define a phenotype for each haplotype within an individual. Therefore, an individual has two phenotypes rather than one.

### 2.3 Scoring localization and detection

To address the question of localization, we scored the distance of the peak association signal from the risk region based on the average absolute value of the distance of peak signals across the entire genomic region. The average distance was used when there were multiple peaks with the same maximum strength of association. Specifically, for each method, on each dataset, we computed the average distance (in bases) of the peak association signals from the risk region and plotted the empirical cumulative distribution function (ECDF) of the average based on the 200 simulated samples. Thus, the ECDF at point *x* is the proportion of the 200 simulated samples with average distance less than or equal to *x*. A method with higher ECDF than another method localizes the signal better.

To detect association with a given method, we used a maximum score across all the SNVs in a dataset to obtain a global test of association across the entire genomic region. We determined the null distribution of the global test statistic for each method by permuting the case-control labels. For the global test statistic, we used either a maximum statistic or the maximum of *−log*_10_ of p-values across the genomic region. The global test statistics for the different association methods are not comparable since they are not on the same scale. To make these statistics comparable across methods, we considered their permutation p-values. We defined these p-values as the proportion of test statistics under the permutation-null distribution that are greater than or equal to the observed value. We then compared the distribution of the resulting p-values for the different methods by plotting their ECDFs.

## 3 Results

In this section, we first present the summaries of our example dataset and the resulting plots from the selected association methods. We then present our results from the simulation study for localizing and detecting the association signal.

### 3.1 Example dataset

#### 3.1.1 Population and sample summaries

In the population of 1500 individuals that was simulated for the example data set, we obtained 4000 SNVs, of which 16 were risk SNVs. Of the 4000 SNVs in the population, 2747 were polymorphic in the sample of 50 cases and 50 controls. Of the 16 risk SNVs in the population, 10 were polymorphic in the case-control sample. The linkage disequilibria between the polymorphic risk SNVs was low; all *r*^2^ values were < 0.1 (results not shown). Table 1 summarizes the physical distances, effect sizes of risk SNVs, age of the most recent common ancestors of the derived alleles in generations, number of recombinations and minor allele frequencies (MAFs) of the 10 risk SNVs in the sample. Of these 10 risk SNVs, the fourth is the oldest but the seventh is the most frequent, owing to the neutral random variation of the simulated trees.

**Table 1:** Summaries for risk SNVs.

| Risk SNV | Position (kbp) | Effect size<sup>a</sup> | Age<sup>b</sup> | NR<sup>c</sup> | MAF |  |  |
| --- | --- | --- | --- | --- | --- | --- | --- |
|  |  |  |  |  | Population | Cases | Controls |
| 1 | 975.0 | 0.039 | 298,737 | NA | 0.001 | 0.01 | 0.00 |
| 2 | 990.0 | 0.079 | 33,0135 | 18 | 0.009 | 0.02 | 0.00 |
| 3 | 990.5 | 0.307 | 802,458 | 0 | 0.013 | 0.07 | 0.01 |
| 4 | 991.5 | 0.380 | 20,690,057 | 2 | 0.039 | 0.07 | 0.03 |
| 5 | 993.0 | 0.380 | 1,498,566 | 0 | 0.031 | 0.08 | 0.02 |
| 6 | 997.5 | 0.156 | 438,663 | 0 | 0.006 | 0.02 | 0.02 |
| 7 | 1005.5 | 1.128 | 3,655,347 | 10 | 0.115 | 0.27 | 0.07 |
| 8 | 1012.5 | 0.232 | 2,115,335 | 5 | 0.013 | 0.05 | 0.01 |
| 9 | 1019.0 | 0.039 | 703,023 | 4 | 0.001 | 0.01 | 0.00 |
| 10 | 1038.5 | 0.195 | 405,567 | 14 | 0.013 | 0.04 | 0.01 |

<sup>a</sup> Effect sizes were computed from MAF in cases and controls and the regression effect, as described in [31].
<sup>b</sup> Age was measured by the number of generations back to the most recent common ancestor of the derived alleles.
<sup>c</sup> NR, the number of recombination events between the current and previous risk SNV in the sample.

Figure 1 compares the distribution of risk haplotypes in cases and controls. We define a risk haplotype to be a haplotype that carries a risk SNV. Figure 2 shows the effect size of the polymorphic risk SNVs versus their location in the risk region (panel (a)) and their age in generations, in the log-base-10 scale, respectively.

**Figure 1:**
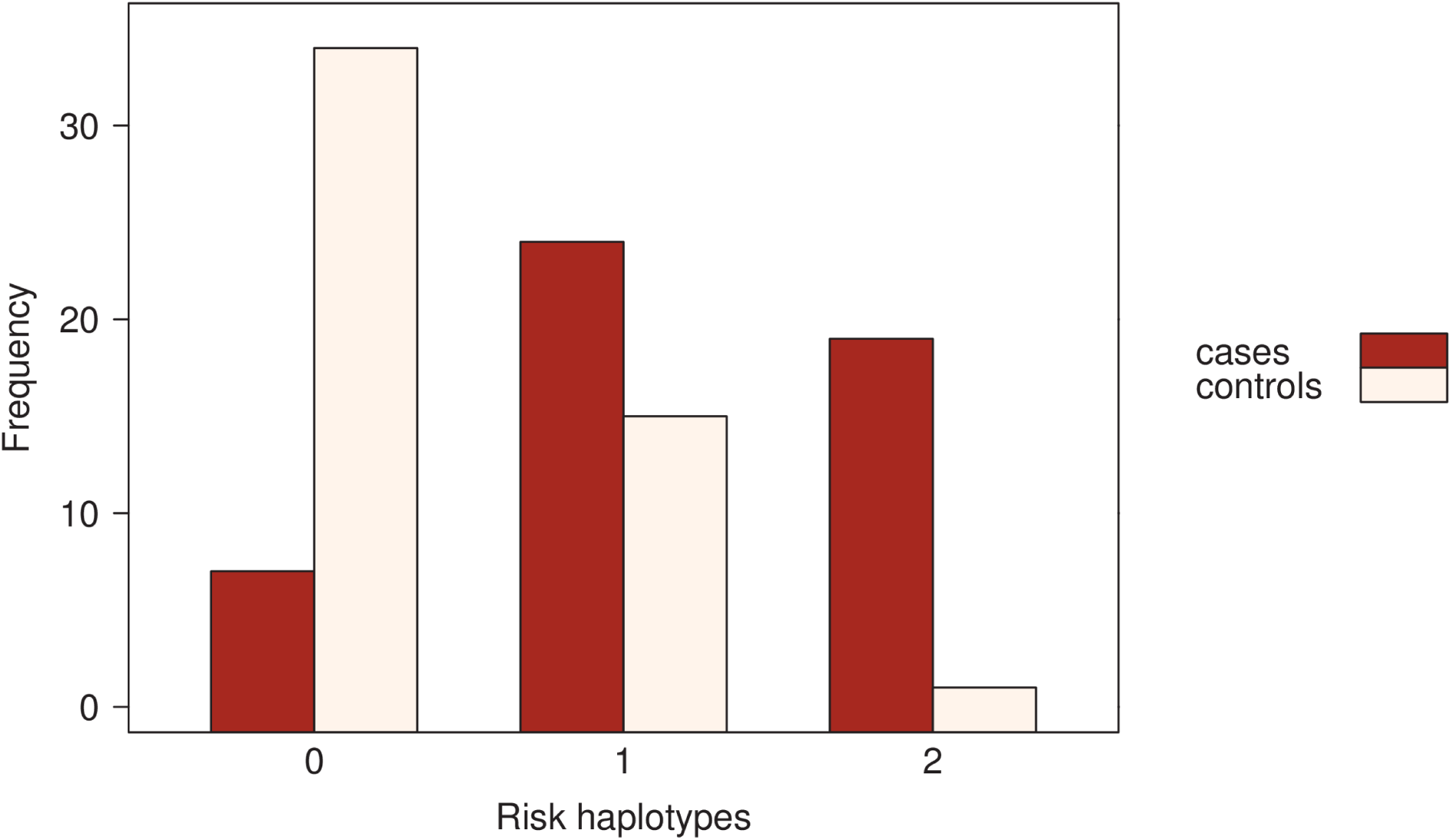
Number of haplotypes that carry risk variants for both cases and controls.

**Figure 2:**
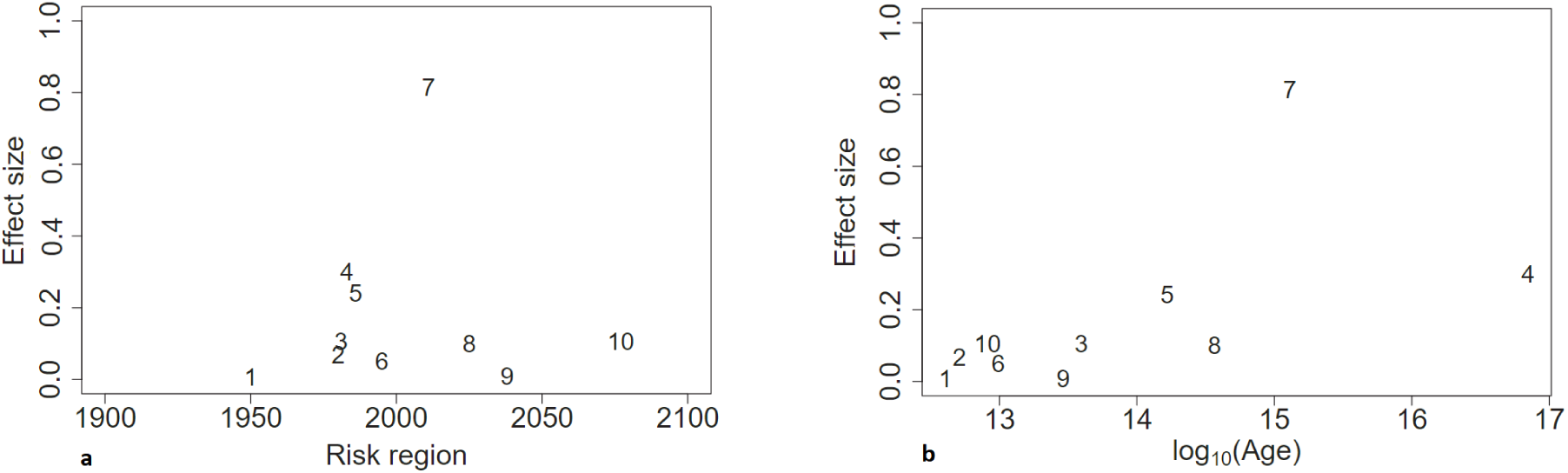
**a)** True effect size of polymorphic risk SNVs versus their positions in the risk region. **b)** True effect size of polymorphic risk SNVs versus their age in generations, in the log-base-10 scale. The risk SNVs are numbered according to their physical location in the risk region.

#### 3.1.2 Association results

Figure 3 shows the resulting plots for each association method using the example dataset. Results from the single-variant method of Fisher’s exact test are shown in panel (a). In our example dataset, Fisher’s exact test does not localize the peak signal, which is distal to the disease-risk region.

**Figure 3:**
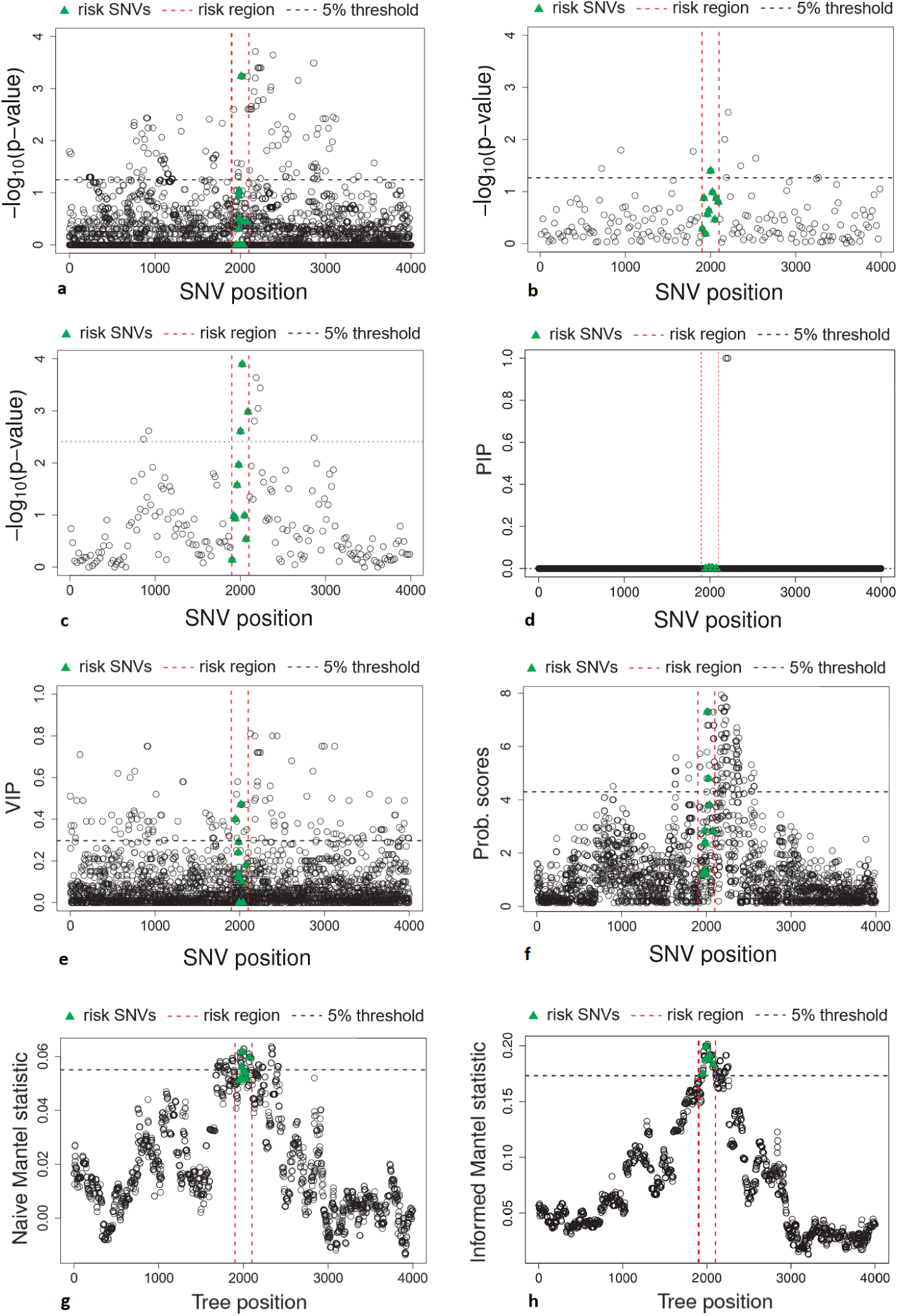
Plots of association results from the eight selected association methods in the 50 cases and 50 controls. **a)** Fisher’s exact test. **b)** VT test. **c)** C-alpha test. **d)** Posterior inclusion probabilities (PIPs) computed from CAVIARBF. **e)** Variable-inclusion probabilities (VIPs) for SNVs computed from elastic net. **f)** Clustering scores for each SNV, using the probability scores criterion in Blossoc. **g)** Naive-Mantel statistics for each tree position (SNV). **h)** Informed-Mantel statistics for each tree position (SNV). The horizontal dashed line represents the 5% significant threshold based on permutation and adjusted for multiple testing across the entire genomic region.

Panel (b) and (c) of Figure 3 show the results from the pooled-variant methods. The C-alpha test in panel (c) has stronger associations than the VT test in panel (b), and the C-alpha test localizes the peak association signal to the disease-risk region whereas the VT test doesn’t.

Panel (d) and (e) of Figure 3 show the results from the joint-modeling methods. The estimated posterior (PIP) and variable (VIP) inclusion probabilities for the SNVs were computed from CAVIARBF (panel (d)) and elastic net (panel (e)), respectively. We used 100 bootstrap samples to estimate VIPs via elastic net. CAVIARBF provides estimates of the PIPs at each SNV. In our example dataset, both elastic net and CAVIARBF show peak signal outside the risk region, but CAVIARBF localizes the signal better than elastic net.

Panel (f), (g) and (h) of Figure 3 show the results from the tree-based methods: Blossoc and the two versions of the Mantel test, i.e. naive- and informed-Mantel. We applied Blossoc to the phased haplotypes, using the probability score criterion for each SNV across the region (panel (f)). In our example dataset, Blossoc shows relatively high association, but the peak signal is outside the risk region. Panel (g) shows the statistics computed from the naive-Mantel test. Our example dataset shows relatively high association signal within the risk region but the peak signal is outside of it. Panel (h) shows the statistics computed from the informed-Mantel test. This informed-Mantel test successfully localizes the peak signal to the risk region.

### 3.2 Simulation study

We first present the simulation results for localizing the association signal, followed by the results for detecting the association signal.

#### 3.2.1 Localizing the association signal

Figure 4 compares the ability of the different methods to localize the association signal. For each of the 200 simulated datasets, we considered the distance of the peak association signal from the risk region. As described in the methods, if there were ties in the peak signal, we took the average distance. The figure shows the ECDFs of these distances for the 200 datasets, for all eight methods. The informed Mantel test outperforms all the other methods; that is, it has the highest proportion of simulated datasets at the lower distance values. Fisher's exact test, C-alpha test, CAVIARBF, and Blossoc perform comparably and relatively well for localizing signal. VT and naive Mantel test have the worst localization performance. As observed in the example dataset, VT has worse performance than C-alpha for localizing the signal.

**Figure 4:**
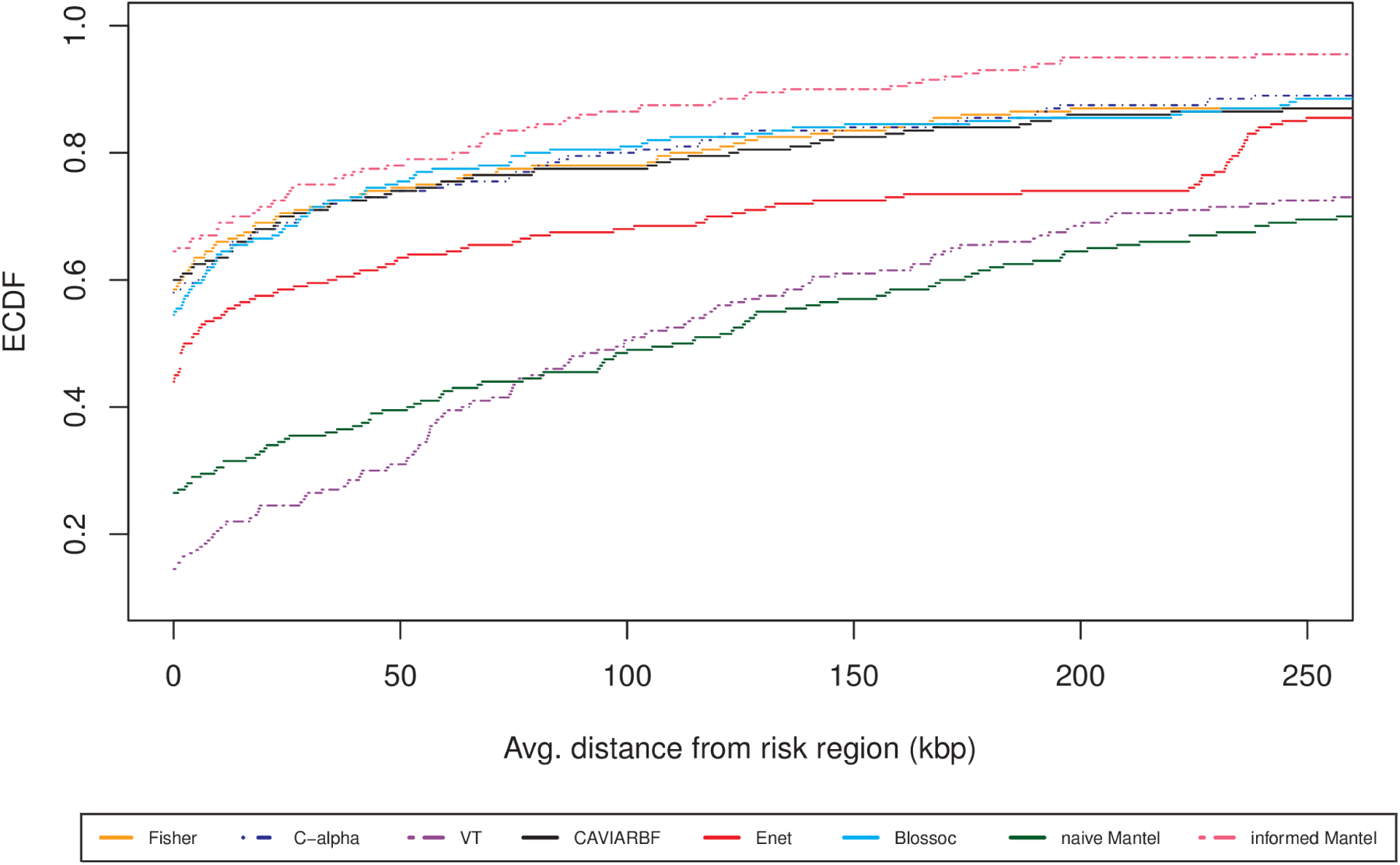
ECDFs of average distances of the peak association signals from the risk region for the 200 datasets. Eight methods are compared: Fisher’s exact test, VT test, C-alpha test, CAVIARBF, Elastic-net, Blossoc, naive Mantel test and informed Mantel test. To better compare methods, the x-axis is shown only for distances ≤ 250kbp.

#### 3.2.2 Detecting the association signal

Figure 5 compares the ability of the methods to detect any association with the disease across the entire genomic region that is being fine-mapped. For each method, we compare the ECDFs, over the 200 datasets, of the permutation p-values computed from the corresponding scores. As expected, the informed Mantel test performs better than all the other methods. The elastic net approach has the lowest power to detect association, followed by the VT approach.

**Figure 5:**
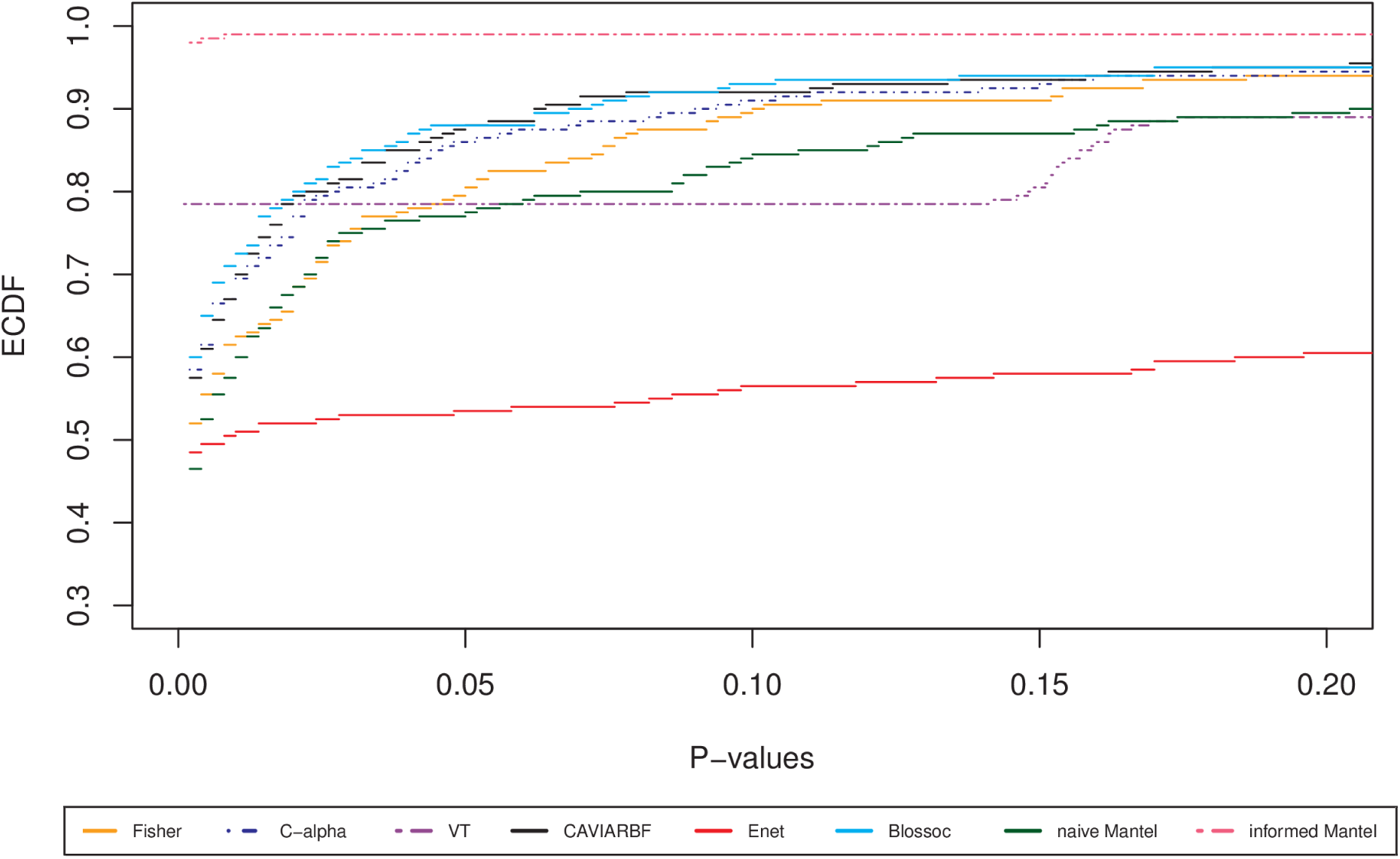
ECDFs of permutation p-values from a global test of association in the genomic region. Eight methods are compared: Fisher’s exact, VT, C-alpha, CAVIARBF, Elastic-net, Blossoc, naive Mantel test and informed Mantel test. The x-axis is shown only for p-values ≤0.20 for a better resolution.

## 4 Discussion

In this study, through coalescent simulation, we have investigated the ability of several popular association methods to fine-map trait-influencing genetic variants in a candidate genomic region. While our simulation investigation can never replace an examination of true sequencing data in the framework where the actual variants are known, it can give some insight into the operating characteristics of these association methods under a popular and tractable model of sequence variation, the coalescent with mutation and recombination. As the first step, we worked through a particular example dataset as a case study for insight into the methods. We then performed a simulation study to score which method localizes the risk subregion most precisely.

In our simulations, the informed Mantel test localized the association signal most precisely among all the methods considered. By contrast, the naive Mantel test performed poorly relative to the other methods. In fact, the naive Mantel test localized the risk region more poorly than Blossoc, CAVIARBF, C-alpha, and Fisher’s exact test. Our results for *localizing* risk variants in *diploid* populations therefore stand in contrast to previous results for *detecting* risk variants in *haploid* populations [5], which found that the naive Mantel test and related tree-based methods performed very well. The poor performance of the naive Mantel test in our simulations can be explained by the misclassification of haplotypes. In haploid populations, case haplotypes without a risk variant are rarer than in diploid populations, and so fewer would be misclassified by the naive case-control phenotypes than in diploid populations. In diploid populations, we do not know which of the two haplotypes in a case carries the disease-risk variant and score both as being “affected”. When only one of the two haplotypes in a case carries a risk variant, defining both haplotypes as “affected” misclassifies one. Therefore, when the majority of cases carry only a single haplotype with risk variants, we expect the informed Mantel test to outperform the naive Mantel test because it defines only case haplotypes that carry the risk variant as “affected”. The development of methods to identify which haplotypes carry risk variants would thus be an avenue for further research in genealogy-based approaches to fine-mapping risk variants.

When computing the naive and informed Mantel statistics, we have assumed that the true genealogical trees are known. In practice, however, these trees are not known. The accuracy of trees reconstructed from the sequence data is expected to affect the localization abilities of the Mantel statistics. When the true trees can be reconstructed with a high degree of accuracy from the available sequence data, the informed Mantel test applied to the reconstructed tree should localize the association signal well. To reconstruct the haplotype partitions implied by the genealogical trees, we applied the methods outlined in [8] to the sequence data. To gain insight into the accuracy of the reconstruction, we computed the Rand index [32] between the true and reconstructed partitions at the genomic position of each risk SNV. The Rand index is a measure between 0 and 1 reflecting the agreement of the partitions. For the ten risk SNVs labelled 1-10 in Table 1, the Rand index values based on ten clusters from each partition were 0.849, 0.814, 0.914, 0.900, 0.900, 0.900, 0.853, 0.885, 0.895, 0.882, respectively. These high values suggest good agreement and accuracy of reconstruction. However, candidate genomic regions with lower mutation and/or higher recombination rates than the rates we have used in our simulations would be expected to have less accurate reconstructions. In these cases, the performance of the Mantel procedures would be expected to be poorer than shown here. The nature and extent of this performance loss would be an interesting topic for future work.

Our simulation study also provides a comparison of the VT to the C-alpha test. Even though the effects are one directional, C-alpha showed higher localization signal in the risk region than VT. Our findings for localization with the VT and C-alpha tests in a diploid population are consistent with those of Burkett et al. [5] for detection in a haploid population. We would in fact expect better performance of the VT test than the C-alpha test since VT is for rare variants having the same direction of effect, which we simulated [14]. However, variance-component tests such as C-alpha have higher power than burden tests such as VT when the proportion of risk variants in the set of tested variants is low [33]. In our example dataset, the highest proportion of risk variants within moving windows of 20 SNVs was 20%. Therefore, a possible explanation for the better performance of C-alpha relative to VT is a relatively low proportion of risk variants within the moving windows. To examine the impact of larger window sizes in our example dataset, we experimented with windows of size 50 and 100 SNVs (overlapping by 5 SNVs) for both the VT and C-alpha tests. However, we could not see any improvement in localizing the association signal (results not shown).

In the example dataset, most risk variants were rare. Of the 10 risk SNVs that were polymorphic in the sample, four were rare with MAF < 1%, five were low frequency with MAF of 1 — 5%, and one was common with MAF > 5%. In addition, a majority of cases carried a single risk haplotype (see figure 1) and most risk haplotypes contained a single risk SNV. These findings in the example dataset suggest that the results under a dominant model of genetic risk would be similar to our results under an additive model. Under a recessive model of disease risk, we would expect the naive Mantel test to perform as well as the informed Mantel test because both haplotypes in a case would tend to carry risk variants, and so misclassification of haplotypes would be minimized. In the example dataset, we found that the C-alpha test and the informed Mantel test were the only methods that successfully localized the association signal. However, the peak signals from all the other methods (Fisher’s test, VT, CAVIARBF, elastic net, Blossoc and naive-Mantel) were close to the disease-risk region.

There are a number of directions for future work. First, we have focused on a simple model of disease risk, with additive effects, no interactions, and no non-genetic covariates. Simulations with more complex risk models would be an area for further research. In the approaches we have considered, the phenotypes can be adjusted for non-genetic covariates. Second, an examination of true sequencing data with known causal variants would be an interesting future direction once such data resources become more readily available to the public. Finally, for tree-based methods, differentiating between case haplotypes that carry or don’t carry risk SNVs improves localization of the risk region. Preliminary work (not shown) suggests that carrier and non-carrier haplotypes can be differentiated based on their number of positively-associated alleles (at level 5%, uncorrected for multiple testing). In future work, we will pursue this idea with informed-Mantel tests on reconstructed trees.

## 5 Acknowledgements

We thank Kelly Burkett for helpful discussions and the Department of Statistics and Actuarial Science at Simon Fraser University for its generous support. This research was funded in part by the Natural Sciences and Engineering Research Council of Canada.

